# The Oropharynx as a Distinct Colonization Site for *staphylococcus aureus* in the Community

**DOI:** 10.1101/137901

**Authors:** Blake M. Hanson, Ashley E. Kates, Elizabeth Mills, Loreen A. Herwaldt, James C. Torner, Jeffrey D. Dawson, Tara C. Smith

**Affiliations:** The Jackson Laboratory for Genomic Medicine, Farmington, USA; Department of Epidemiology, The University of Iowa, Iowa City, USA; Department of Biostatistics, The University of Iowa, Iowa City, USA; Department of Internal Medicine, The University of Iowa, Iowa City, USA; Department of Biostatistics, Environmental Health Sciences & Epidemiology, Kent State University, Kent, USA

**Keywords:** Staphylococcus aureus, Methicillin-Resistant Staphylococcus aureus, oropharynx, Nasal Cavity, Cohort Studies

## Abstract

**Background:** *S. aureus* is a frequent cause of hospital and community associated infections and colonization is known to increase the risk of infection, with the nares considered the most important colonization site.

**Methods:** We compared the prevalence of nasal and oropharyngeal carriage in a yearlong, prospective cohort study of people from the community as well as assessed risk factors for nares-only and oropharynx-only colonization.

**Results:** Colonization at both anatomical sites was correlated; however, oropharynx only carriage occurred and oropharyngeal swabs were more sensitive than nasal swabs at detecting carriage (77.27% and 72.725 respectively). Non-Caucasian race, having a greater number of people living in your home and more children in your home all significantly increased the odds of oropharynx-only carriage. Having *S. aureus* present on home environmental sites, exercising in a fitness center, and sharing bath towels all increased the odds of nares-only carriage.

**Conclusions:** Oropharyngeal swabs increase the detection of *S. aureus* colonization in community embers.

## INTRODUCTION

*S. aureus* is a commensal, gram-positive bacterium colonizing about 30% of the population [1], causing a wide range of infections. The anterior nares (nostrils) have long been considered the most important site of colonization [2, 3]. While *S. aureus* colonization may not be harmful to the host, it increases the risk of symptomatic infections [4]. Persons colonized with *S. aureus* are asymptomatic carriers able to transmit the bacterium to susceptible persons via direct transmission or indirect contact with fomites [5].

*S. aureus,* specifically methicillin-resistant *Staphylococcus aureus* (MRSA), initially emerged in the 1960s as a hospital-associated pathogen [6–8], and has been observed to cause a wide variety of healthcare-associated infections from skin and soft-tissue infections (SSTIs) to osteomyelitis and sepsis [9]. In the mid-1990s, however, MRSA infections began emerging in populations who had no known exposure to hospitals or healthcare systems [10].

Recent emerging trends in antimicrobial resistance in *S. aureus* [11] have prompted researchers and clinicians to shift their focus from treatment to prevention. *S. aureus* is often resistant to numerous antimicrobial agents and resistance mechanisms transfer readily [12]. Studies assessing the effectiveness of nasal decolonization have shown 56% of persons decolonized with mupirocin and 52% of persons decolonized with chlorhexidine [13] became recolonized, with many decolonized persons reacquiring their original colonizing strain [14], possibly because they were colonized at a separate anatomical location not affected by the decolonization regimen or because their environment was contaminated.

Previous studies have assessed rates of extra-nasal colonization at sites such as the axilla, perineum, inguinal folds [15–18], and the pharynx [16, 19]. Most previously published studies focused on persons receiving healthcare or persons with a history of prior *S. aureus* infections. Within these populations, the prevalence of oropharyngeal colonization ranged from 5.8% to 30% [15, 17, 20–22]. While there are few reports on oropharyngeal colonization rates in the community, studies have shown oropharyngeal colonization rates range from 10.8% to 46.5% [23, 24], which are equal to or greater than rates observed in the healthcare setting. In fact, Mertz et al. hypothesized oropharyngeal *S. aureus* carriage may be more common, and thus more important, in the community than in the healthcare setting [19].

Because few studies have assessed oropharyngeal colonization in healthy persons and because a high percentage of decolonized patients become recolonized, further research assessing the importance of oropharyngeal colonization in healthy persons is critical. In this study, we assess whether the oropharynx is an important site for *S. aureus* colonization as well as colonization risk factors in a population of healthy community members.

## METHODS

### Study design and sample collection

Participants from 95 families were enrolled in a prospective cohort study from two counties in Iowa, Johnson County and Keokuk County. Enrollment occurred between October 2011 and January 2012 with participants being followed weekly for 52 weeks. The methodologies of enrollment, sample collection, and molecular characterization have been described previously. Briefly, participants were enrolled in their homes by the research team. The University of Iowa institutional review board approved all study protocols prior to recruitment. At enrollment, the research team swabbed six commonly touched locations in the home (primary television remote, main bathroom toilet flush lever and light switch, kitchen sink handle, refrigerator door handle, and oven knobs) for *S. aureus* contamination. Participants were also trained on how to self-swab their nares (adults and minors) and their oropharynx (adults). Participants then swabbed themselves and mailed their samples to the research team once a week for 52 weeks.

### Statistical analysis

Data were analyzed using SAS software version 9.3 (Cary, NC, USA) and R version 3.3.1. Participant demographics were compared between the two counties, using the Student’s T-Test to assess age, the Chi-Squared Test without Yate’s Continuity Correction to assess gender, and the Fisher’s Exact Test to assess race, given the small number of non-Caucasian participants.

Colonization status was determined by dividing the total number of positive cultures by the total number of nasal and oropharyngeal cultures obtained during the study. Participants were classified as: non-carriers if <50% of cultures from both the nares and the oropharynx were positive; oropharynx-only carriers if ≥50% of oropharyngeal cultures were positive, but <50% of the nares cultures were positive; nares-only carriers if ≥50% of nares cultures were positive but <50% of oropharyngeal cultures were positive; and nasal and oropharyngeal carriers if ≥50% of cultures from both sites were positive. These cutoffs were used to define equal quartiles within the data for risk factor analyses. Sensitivity of the anatomical sites was calculated by dividing the total number of a subject’s positive samples at the given site (totaling those positive at both sites and those only positive for the given site) by all colonized persons. Confidence intervals for the sensitivities were done using exact binomial confidence limits.

Logistic regression with receiver operating curves (ROC) was used to determine the lowest number nares and oropharynx samples in tandem necessary to accurately predict the carrier status of each participant. Carrier status was used as the dependent variable; the number of positive nares cultures out of the total number obtained (i.e. 4 positive of 10 cultures) and the number of positive oropharyngeal cultures out of the total cultures obtained were used as independent variables. The ROC analysis was used to calculate false positive rates (FPR) and true positive rates (TPR).

Random effects logistic regression with Satterthwaite’s approximation for degrees for the degrees of freedom was used to assess risk factors for both oropharyngeal and nasal colonization with a p-value ≤0.05 considered significant. This method was used to account for the random effects of both family and county. All risk factors assessed were modeled as dichotomous or categorical variables, except for the number of children in the family and house size, which were modeled as continuous variables. After adjustment for individual risk factors and the random effect of family in the bivariate analysis, some models had a non-positive variance for county, indicating the remaining variance was not attributable to county, and thus the variance for county was set to zero [25]. If cell counts for an analysis included a zero, the Fisher’s Exact Test was used to assess the association between the risk factor and being a carrier. This method is unable to account for the random effects. The model assessing oropharynx-only carriage was adjusted for age.

The Pearson product-moment correlation coefficient (PPMC) was used to determine the nature of the relationship between nasal and oropharyngeal colonization. PPMC was also used to determine differences in the relationship of nasal and oropharyngeal colonization by county. A regression line with 95% confidence intervals were plotted for each county.

## RESULTS

One hundred and seventy-seven adults and 86 minors from 95 families were enrolled into the study. As minors submitted only nares swabs, they were excluded from these analyses. The average age of the adult participants was 44.5 (range: 23.1-67.7 years). Three participants did not indicate their age. Of the 177 participants, 55.2% were female; 2 participants did not indicate their gender. Most (92.2%) participants considered themselves Caucasian, with 7.8% of participants indicating “other” (American Indian/Alaska native, Asian, Hispanic or Latino, or Native Hawaiian or other Pacific Islander) as their race.

A logistic regression model with ROC curves found that 14 consecutive swabs was the minimum number of swabs necessary to minimize the FPR, maximize the TPR, and minimize the number of swabs obtained. Using 14 consecutive swabs, we observed an FPR of 0.0763 and a TPR of 0.7586 (Table 1, Figure 1). Of the 177 adults, 152 (85.9%) submitted 14 or more swabs, with an average of 34.7 swab sets per participant (range: 14-52), and were included in further analyses. One hundred and twenty-nine participants of the 152 (84.9%) had at least one swab positive for *S. aureus*. One hundred and eight (71.1%) participants were classified as non-carriers, 10 (6.6%) were nares-only carriers, 12 (7.9%) were oropharynx-only carriers, and 22 (14.5%) were nares and oropharynx carriers (Figure 2). Additional County and gender specific information can be found in Table 2. Oropharyngeal swabs had greater sensitivity compared to oropharyngeal swabs at 72.7% and 77.3%, respectively (Table 3).

**Table 1.** ROC curve descriptive statistics at multiple informative levels.

| **Number of Timepoints** | **True Positive** | **True Negative** | **False Positive** | **False Negative** | **Total<sup>\*</sup>** | **TPR** | **FPR** |
| --- | --- | --- | --- | --- | --- | --- | --- |
| **1** | 18 | 196 | 11 | 12 | 237 | 0.6297 | 0.1017 |
| **7** | 20 | 107 | 9 | 11 | 147 | 0.6896 | 0.0932 |
| **10** | 10 | 108 | 9 | 20 | 147 | 0.6896 | 0.0847 |
| **14** | 22 | 109 | 7 | 9 | 147 | 0.7586 | 0.0763 |
| **19** | 23 | 104 | 6 | 8 | 141 | 0.7931 | 0.0714 |

<sup>\*</sup>Sample size at each time point decreases due to participant attrition over the sampling period.

**Table 2.** Distribution of participants’ carrier status by county and gender

|  | **Non-carrier** | **Nares-only carrier** | **Oropharynx-only carrier** | **Nasal and Oropharyngeal carrier** |
| --- | --- | --- | --- | --- |
| County |  |  |  |  |
| Johnson (n=73) | 49 (67.1%) | 1 (1.4%) | 11 (15.1%) | 12 (16.4%) |
| Keokuk (n=79) | 59 (74.7%) | 9 (11.4%) | 1 (1.3%) | 10 (12.7%) |
| Gender |  |  |  |  |
| Male (n=66) | 43 (65.2%) | 5 (7.6%) | 5 (7.6%) | 13 (19.7%) |
| Female (n=84) | 63 (75.0%) | 5 (6.0%) | 7 (8.3%) | 9 (10.7%) |
| Total (n=152) | 108 (71.1%) | 10 (6.6%) | 12 (7.9%) | 22 (14.5%) |

**Table 3.**
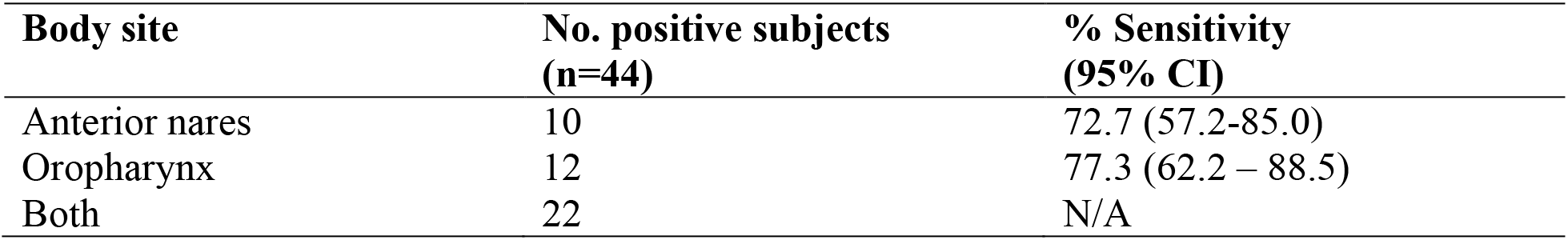
Sensitivity of the swabbing sites for detection of *S. aureus*

| **Body site** | **No. positive subjects**<br>**(n=44)** | **% Sensitivity**<br>**(95% CI)** |
| --- | --- | --- |
| Anterior nares | 10 | 72.7 (57.2-85.0) |
| Oropharynx | 12 | 77.3 (62.2 – 88.5) |
| Both | 22 | N/A |

**Figure 1.**
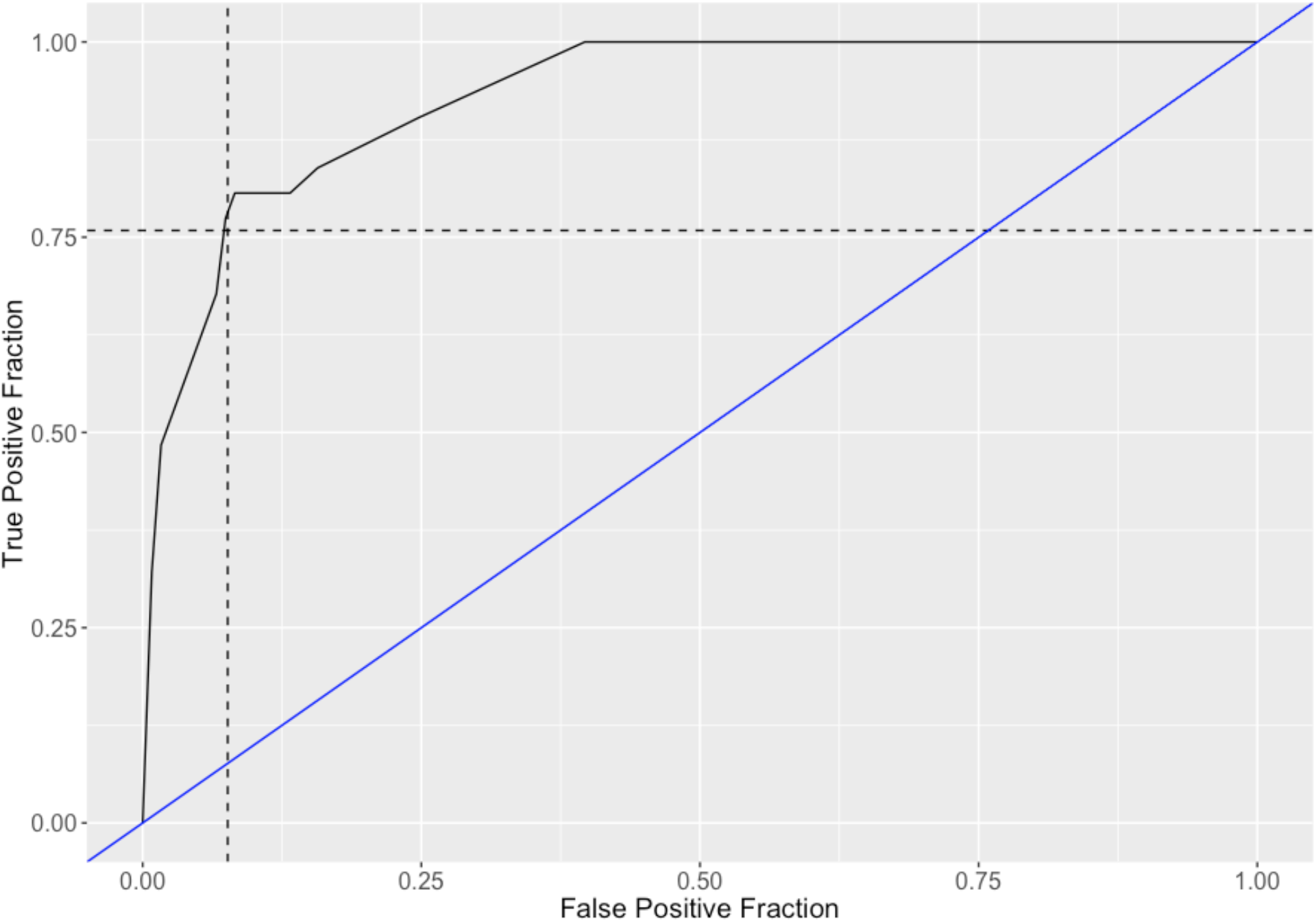
ROC curve of the number of swabs required to differentiate S. aureus carrier states.

**Figure 2.**
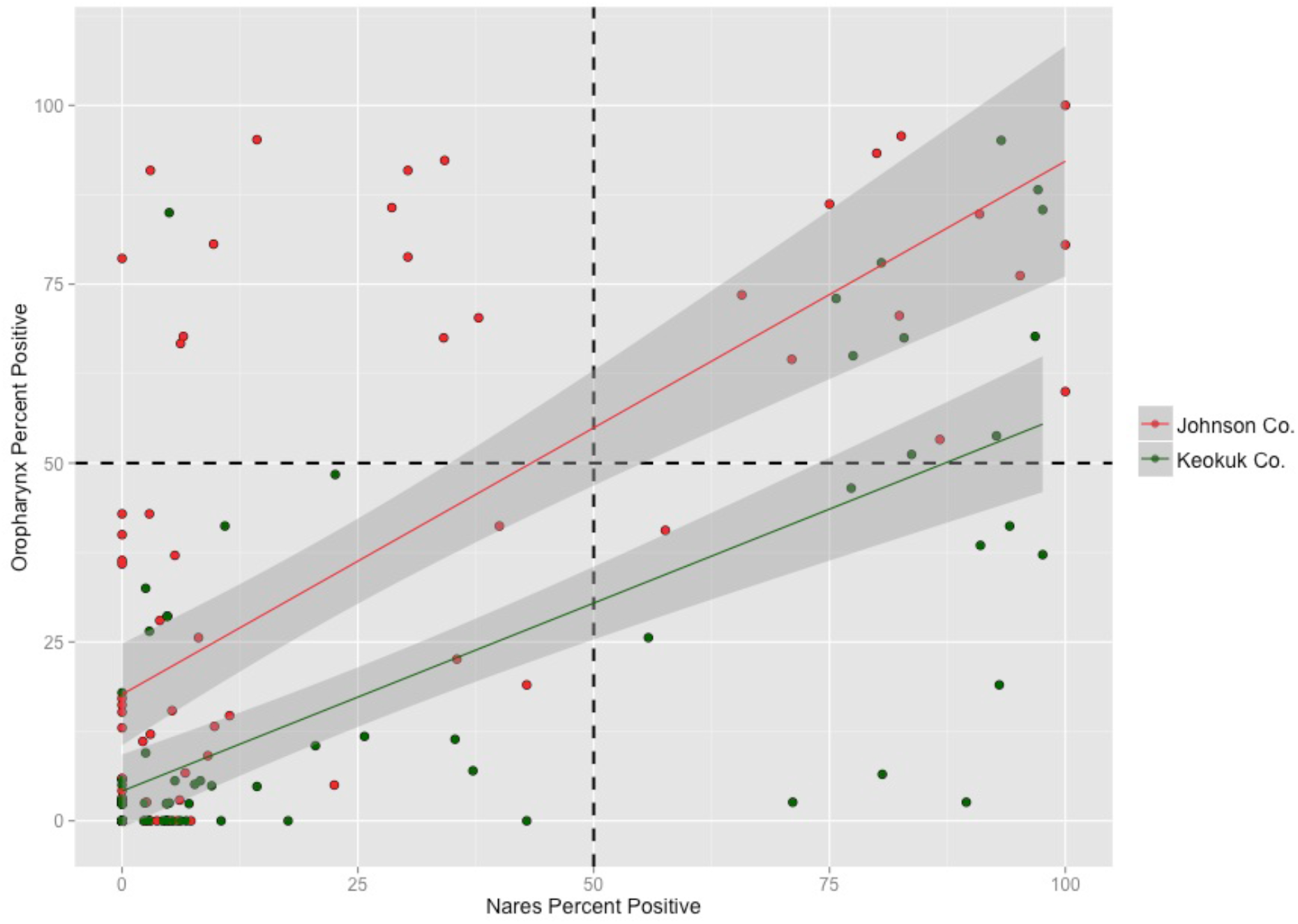
Scatterplot with regression lines and 95% confidence intervals showing the relationship between oropharyngeal and nasal colonization by county. Each circle denotes a single participant. Hash lines indicatethe 50% cut-offs used to separate the data into quadrants. The top right quadrant represents nasal and oropharyngeal carriers, top left represents oropharyngeal-only carriers, bottom left represents non-carriers, and the bottom right represents nares-only colonizers.

Random effects logistic regression analyses were used to identify significant risk factors for oropharynx-only and nares-only colonization. We identified significant relationships between oropharynx-only colonization and race (OR: 8.8, 95% CI: 1.49, 52.4, p-value = 0.017), with participants of non-Caucasian race having increased odds of colonization. House size (OR: 1.86, 95% CI: 1.08, 3.20, p-value = 0.027) and number of children within the household (OR: 1.98, 95% CI: 1.14, 3.43, p-value = 0.016) also increased the odds of oropharyngeal-only colonization as the number of people as well as the number of children in the home increased (Table 4).

**Table 4.** Risk for oropharyngeal colonization compared to non-carriers

| **Risk Factor** | **Oropharyngeal Carrier (%)** | **Total** | **OR (95% CI)** | **P-value** |
| --- | --- | --- | --- | --- |
| Age | 108 (90.0%) | 120 | 0.99 (0.93, 1.06) | 0.7572 |
| Gender |  |  |  |  |
| Male | 62 (89.9%) | 69 |  |  |
| Female | 44 (89.8%) | 49 | 0.92 (0.25, 3.35) | 0.9001 |
| Race |  |  |  |  |
| Caucasian | 103 (92.8%) | 111 |  |  |
| Other | 5 (55.6%) | 9 | 8.84 (1.49, 52.4) | 0.0169 |
| House Size |  |  |  |  |
| One | 1 (100%) | 1 |  |  |
| Two | 51 (92.7%) | 55 |  |  |
| Three | 30 (100%) | 30 |  |  |
| Four | 15 (79.0%) | 19 |  |  |
| Five | 10 (76.9%) | 13 |  |  |
| Six | 0 (NA) | 0 |  |  |
| Seven | 1 (50.0%) | 2 | 1.86 (1.08, 3.20) | 0.0273 |
| Number of children in the home |  |  |  |  |
| Zero | 60 (95.2%) | 63 |  |  |
| One | 20 (95.2%) | 21 |  |  |
| Two | 19 (82.6%) | 23 |  |  |
| Three | 8 (72.3%) | 11 |  |  |
| Four | 0 (NA) | 0 |  |  |
| Five | 1 (50.0%) | 2 | 1.98 (1.14, 3.43) | 0.0164 |
| Number of *S. aureus* positive environmental sites |  |  |  |  |
| Zero | 82 (89.1%) | 92 |  |  |
| One | 15 (88.2%) | 17 |  |  |
| Two | 9 (100%) | 9 |  |  |
| Three | 1 (100%) | 1 |  |  |
| Four | 0 (NA) | 0 |  |  |
| Five | 1 (100%) | 1 | 0.55 (0.14, 2.14) | 0.3862 |
| Asthma |  |  |  |  |
| No | 99 (89.2%) | 111 |  |  |
| Yes | 8 (100%) | 8 | NA | 0.5971 |
| Eczema |  |  |  |  |
| No | 98 (89.1%) | 110 |  |  |
| Yes | 10 (100%) | 10 | NA | 0.5979 |
| Other Skin Conditions |  |  |  |  |
| No | 98 (89.9%) | 109 |  |  |
| Yes | 10 (90.9%) | 11 | 0.97 (0.09, 0.39) | 0.9820 |
| Diabetes |  |  |  |  |
| No | 103 (90.4%) | 114 |  |  |
| Yes | 5 (83.3%) | 6 | 0.36 (0.02, 5.78) | 0.4659 |

Table 4: Risk for oropharyngeal colonization compared to non-carriers continued
| **Risk Factor** | **Oropharyngeal Carrier (%)** | **Total** | **OR (95% CI)** | **P-value** |
| --- | --- | --- | --- | --- |
| Disorder of the immune system |  |  |  |  |
| No | 97 (89.0%) | 109 |  |  |
| Yes | 6 (100%) | 6 | NA | 1.0 |
| Heart condition |  |  |  |  |
| No | 103 (89.6%) | 115 |  |  |
| Yes | 5 (100%) | 5 | NA | 1.0 |
| Cancer |  |  |  |  |
| No | 100 (89.3%) | 112 |  |  |
| Yes | 8 (100%) | 8 | NA | 0.5977 |
| Cancer medications |  |  |  |  |
| No | 105 (89.7%) | 117 |  |  |
| Yes | 3 (100%) | 3 | NA | 1.0 |
| Used an antibiotic in the last 90 days |  |  |  |  |
| No | 95 (89.6%) | 106 |  |  |
| Yes | 13 (92.9%) | 14 | 1.25 (0.12, 13.51) | 0.6794 |
| Family member had an SSTI |  |  |  |  |
| No | 98 (89.1%) | 110 |  |  |
| Yes | 7 (100%) | 7 | NA | 0.5960 |
| Participant had an SSTI |  |  |  |  |
| No | 106 (89.8%) | 118 |  |  |
| Yes | 2 (100%) | 2 | NA | 1.0 |
| Participant worked in healthcare |  |  |  |  |
| No | 85 (91.4%) | 93 |  |  |
| Yes | 21 (84.0%) | 25 | 0.75 (0.17, 3.27) | 0.6969 |
| Visited a hospital in last 90 days. |  |  |  |  |
| No | 63 (87.5%) | 72 |  |  |
| Yes | 44 (93.6%) | 47 | 2.04 (0.43, 9.73) | 0.2479 |
| Admitted to an outpatient facility |  |  |  |  |
| No | 99 (89.2%) | 111 |  |  |
| Yes | 7 (100%) | 7 | NA | 1.0 |
| Family member hospitalized in last 90 days |  |  |  |  |
| No | 105 (90.5%) | 116 |  |  |
| Yes | 3 (75%) | 4 | 0.33 (0.02, 5.87) | 0.3666 |

Table 4: Risk for oropharyngeal colonization compared to non-carriers continued
| **Risk Factor** | **Oropharyngeal Carrier (%)** | **Total** | **OR (95% CI)** | **P-value** |
| --- | --- | --- | --- | --- |
| Had a child in daycare |  |  |  |  |
| No | 89 (89.0%) | 100 |  |  |
| Yes | 19 (95.0%) | 20 | 3.71 (0.38, 36.12) | 0.2566 |
| Played team sports |  |  |  |  |
| No | 103 (90.4%) | 114 |  |  |
| Yes | 4 (80.0%) | 5 | 0.59 (0.05, 7.58) | 0.6851 |
| Had a family member play team sports |  |  |  |  |
| No | 69 (94.5%) | 73 |  |  |
| Yes | 32 (82.1%) | 39 | 0.17 (0.03, 0.97) | 0.0464 |
| Used a fitness center |  |  |  |  |
| No | 74 (91.4%) | 81 |  |  |
| Yes | 31 (86.1%) | 36 | 1.25 (0.30, 5.32) | 0.7565 |
| Shared bath towels with other members of the household |  |  |  |  |
| No | 67 (90.5%) | 74 |  |  |
| Yes | 41 (89.1%) | 46 | 1.23 (0.29, 5.13) | 0.7747 |
| Shared hand towels with other members of the household |  |  |  |  |
| No | 16 (88.9%) | 18 |  |  |
| Yes | 92 (90.2%) | 102 | 1.45 (0.21, 9.95) | 0.7030 |
| Used antibacterial hand soap |  |  |  |  |
| No | 25 (86.2%) | 29 |  |  |
| Yes | 83 (91.2%) | 91 | 0.92 (0.20, 4.21) | 0.9152 |
| Participant used tobacco products |  |  |  |  |
| No | 100 (90.1%) | 111 |  |  |
| Yes | 7 (87.5%) | 8 | 0.78 (0.07, 9.13) | 0.8402 |

Several risk factors for nares-only colonization were identified. Having *S. aureus* positive environmental sites within the home increased the odds of nares-only colonization (OR: 2.02, 95% CI: 1.11, 3.66, p-value = 0.021), with the odds increasing with the number of *S. aureus* positive sites. We also found using a gym facility/ fitness center (p-value = 0.06) and sharing bath towels with others in the home (p-value = 0.01) increased the odds of nares-only colonization. Participants receiving medication for a cancer diagnosis were significantly less likely to nasally carry *S. aureus* (OR: 0.08, 95% CI: 0.01, 0.69, p-value = 0.022) compared to non-carriers (Table 5).

**Table 5.**
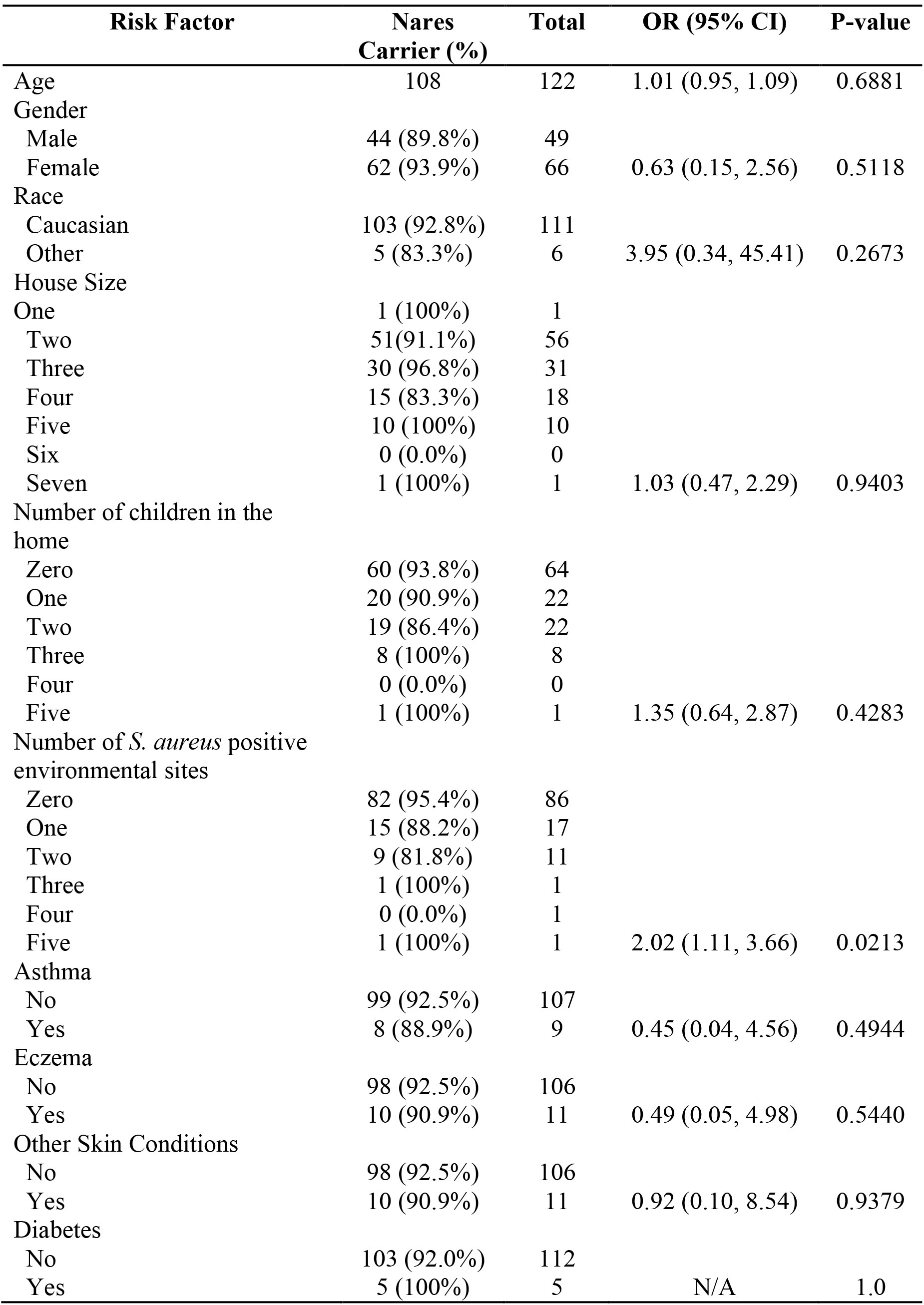
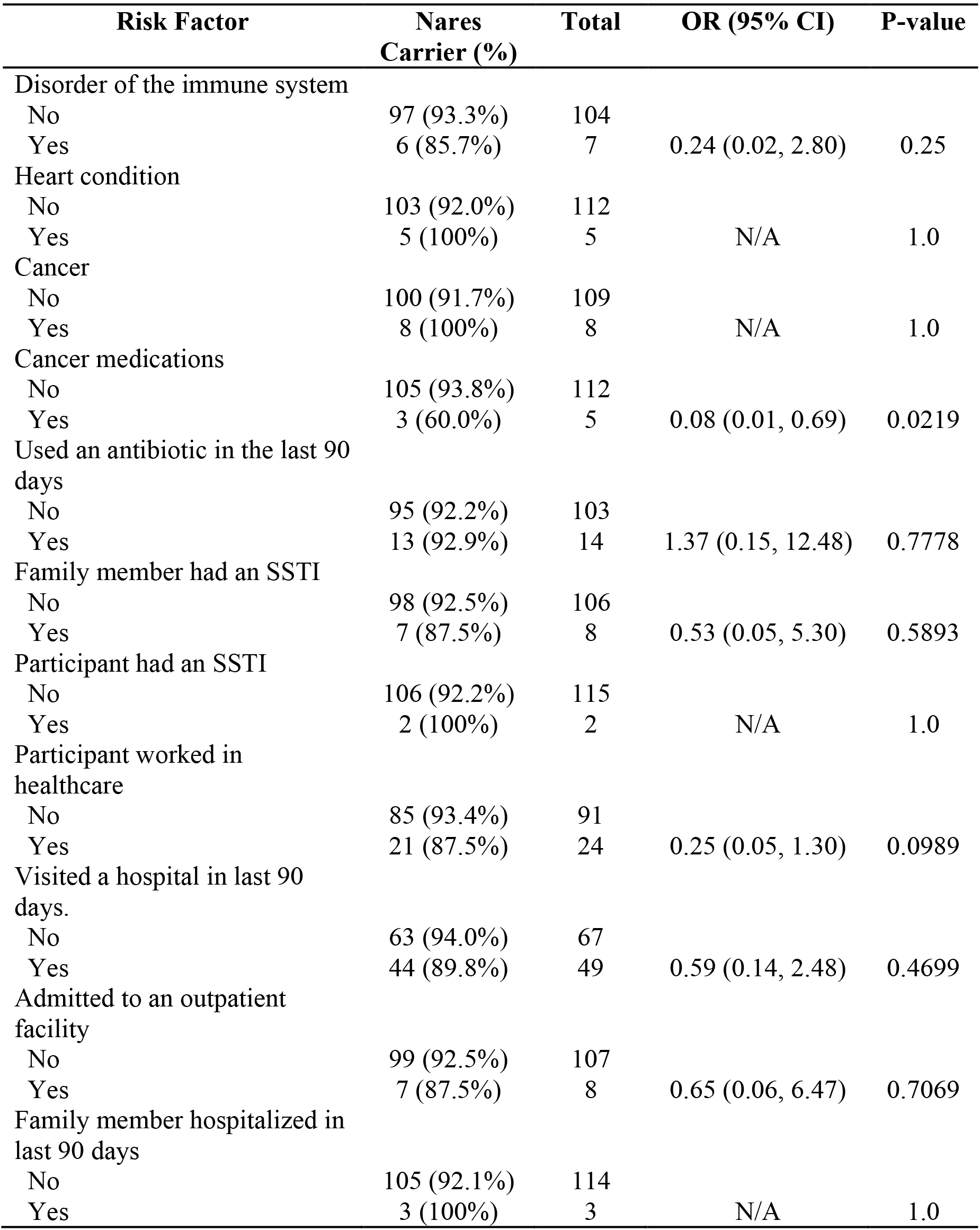
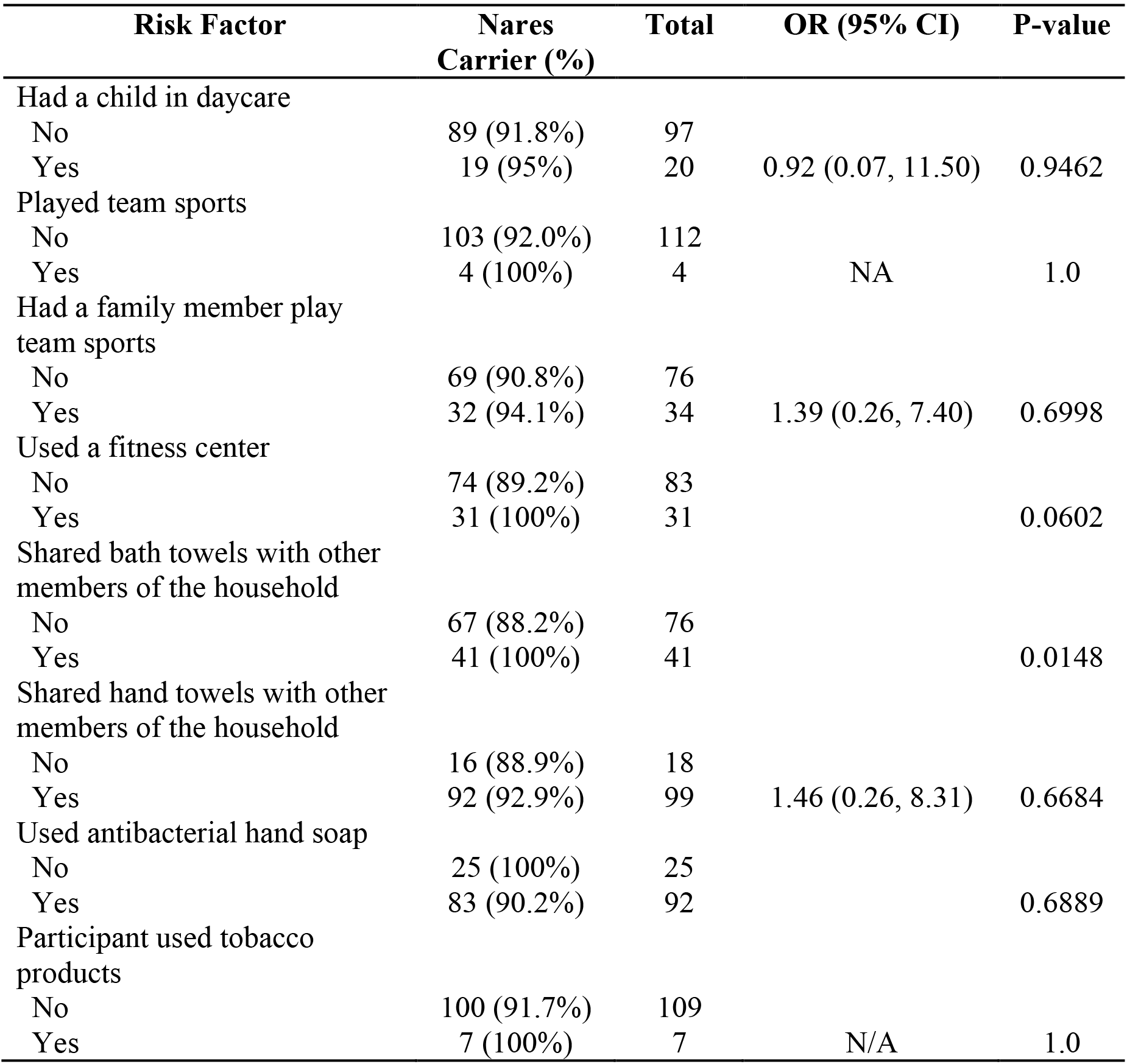
Risk factors for nasal colonization compared to non-carriers

The estimate for the PPMC correlation was 0.64, with a 95% confidence interval of 0.54-0.72 (p-value: <0.001) indicating there is a positive, linear relationship between nasal and oropharyngeal colonization indicating nasal colonization is associated with increased oropharyngeal colonization. We also calculated PPMC by county. The correlation estimate for Johnson County was 0.68 with a 95% confidence interval of 0.53-0.78 and the correlation estimate for Keokuk County was 0.71 with a 95% confidence interval of 0.58-0.81. In addition, the slope of the fitted regression line for Johnson County was 0.75 and the slope for Keokuk County was 0.53. The 95% confidence intervals for each county did not overlap at any point (Figure 2). The results of the Pearson chi-squared test indicated there was a difference in the sensitivity of the oropharyngeal cultures to detect carriage by county (p-value = 0.0019).

## DISCUSSION

Previous studies that have assessed the prevalence of *S. aureus* colonization in the nares and oropharynx among patients in a healthcare setting and among healthy persons [23, 24, 26, 27], but published studies have not assessed multiple cultures obtained at regular intervals to evaluate the relationship between colonization of these two anatomical sites. Additionally, we found 12 participants (7.89%) who met our definition of *S. aureus* colonization of the oropharynx only, suggesting the oropharynx is likely a distinct colonization site and isolation of *S. aureus* from the oropharynx is not solely due to cross-site contamination from the nares as has been hypothesized previously [28, 29].

Using ROC analysis, we determined 14 consecutive nasal and oropharyngeal swabs were necessary to maximize the TPR and minimize the FPR. For our cohort, the TPR for 14 swabs was 0.793 with and FPR of 0.076. Prior studies have aimed to determine how many swabs are necessary to accurately differentiate between non-carriers, intermittent carriers, and persistent carriers and to establish a “culture rule” to be used for future epidemiologic studies. The importance of accurately differentiating the carriage states has long been understood with persistent carriers having higher bacterial load of *S. aureus* and thus greater potential to shed the bacterium into the environment [30] and a greater risk of subsequent infection [31]. Prior studies have traditionally used 10-12 weeks as to differentiate colonization states [3]. Nouwen et al. aimed to determine the number of swabs necessary to differentiate intermittent colonization from persistent carriers and non-carriers. In their study, they determined only one negative culture was enough to rule out persistent carriage; however, seven weekly swabs were necessary to differentiate intermittent carriage from non-carriers [3]. In our study, seven swabs produced a TPR of .6896 and an FPR of 0.093. There are several differences between our study and the Nouwen study. First, the Nouwen study stopped swabbing at 12 weeks while we have up to 52 weeks of data for ours. Second, the Nouwen study had swabs collected by only one researcher, while which may lead to less misclassification of the carrier state. However, it is not practical in most epidemiologic studies to have one researcher complete all data collection, a limitation recognized by Nouwen et al. [3]. These differences could partially explain the difference in results between the two studies. Fourteen swabs were determined to best minimize the FPR and maximize the TPR in our cohort with increasing the number of swabs not markedly improving either metric. Future studies aiming to differentiate between the carrier states should aim to include 14 weeks of swab collection when feasible.

In our assessment of the relationship between the colonization of the oropharynx and colonization of the nares, the Pearson correlation coefficient of 0.64 indicates colonization at the two anatomical sites is correlated. This correlation implies there is a statistical association between colonization of the two sites and indicates colonization of both the nares and oropharynx is common and related. However, when considering the importance of colonization at each site with regard to capturing all colonized persons, we found oropharyngeal cultures to be more sensitive than nasal cultures (77.3% for the oropharynx vs. 72.7% for the nares). Of the 152 participants, 7.9% were oropharynx-only carriers and 6.6% were nares-only carriers. Several studies have assessed the occurrence of *S. aureus* carriage in the oropharynx and have found it to be an important site for carriage, though these studies have been in healthcare settings [32, 33] or in children [34]. Thus, our results support the addition or oropharyngeal cultures when screening for *S. aureus* colonization, especially in persons presenting from the community.

While the two county-specific PPMC coefficient confidence intervals overlap, a liberal interpretation of the two estimates is oropharyngeal cultures were more likely to increase the yield in Johnson County than in Keokuk County. The results of the Pearson chi-squared test indicated there was a difference in the sensitivity of the oropharyngeal cultures for detecting *S. aureus* carriage rates by county (p-value = 0.0019). These findings support past research that has shown while nasal and oropharyngeal colonization is related, including oropharyngeal swabs greatly increases the sensitivity of identifying all colonizers [33]. The fact the confidence intervals for the county-specific slopes did not overlap further supports this interpretation. Participants from Keokuk County were more likely than participants from Johnson County to carry *S. aureus* in their nares and were also older. Underlying differences in the urban and rural settings not captured in this study might help explain the observed difference in oropharyngeal colonization rates by county.

Several risk factors were identified in relation to both oropharyngeal-only and nasal-only carriage. Non-Caucasian race, greater numbers of people living in the household, and more children in the household all increased the odds of carrying *S. aureus* in the oropharynx; however, none of these were risk factors for nares-only colonization. For the nares, having environmental sites within the home contaminated with *S. aureus* increased the risk of nasal carriage. Working out in a fitness center as well as sharing bath towels with other members of the household also increased the risk of nasal-only colonization. An interesting finding was being on cancer medications was protective against colonization in the nares; however, the number of participants on medication for any cancer was small (n=5), so it is difficult to draw any conclusions. Due to small sample sizes, additional research is needed to further assess risk factors for oropharynx-only and for nares-only *S. aureus* carriage.

We adjusted for familial clustering of *S. aureus* when we assessed risk factors as well as for variation due to population density by enrolling participants from rural and urban counties. The adjustment for these variances improved the accuracy of our risk assessment. Though it is a potential limitation of our study design, participants self-reported risk factor data which may have misclassified some participants due to recall bias. However, this allowed us to assess risk factors that are difficult or impossible to collect via alternative methodologies, such as medical records. We evaluated exposures during the three months before participants enrolled, which improved our ability to identify exposures, as it maximized the timeframe assessed while minimizing recall bias.

Participants in our study provided self-swabbed samples. Lautenbach et al. previously demonstrated patient-collected and provider-collected samples have excellent agreement rates, with participant-collected swabs having greater sensitivity for all anatomic sites tested which included the nose and throat [35]. Delacour et al. demonstrated the transport system we used maintained *S. aureus* viability for 18 days after collection [36]. In addition, we did on-site training to teach our participants and we provided biological sample mailers. Thus, this methodology allowed the participants to collect samples when it was most convenient for them while providing an accurate assessment of *S. aureus* colonization.

Our results suggest additional studies with larger sample sizes are needed to further evaluate the frequency of oropharynx-only *S. aureus* carriage and to identify risk factors for persistent *S. aureus* carriage in the oropharynx without persistent nasal carriage. Further studies are needed to determine why rates of oropharyngeal *S. aureus* carriage vary by rurality. For example, future studies could evaluate the role of environmental contamination and assess exposures, such as livestock contact, in more detail than was possible in this study. We identified healthy persons who carried *S. aureus* only in their oropharynx. This observation strongly supports the hypothesis that the oropharynx is an independent colonization site and supports screening of the oropharynx of healthy adults for *S. aureus* colonization in situations where *S. aureus* screening is warranted. Previous studies have determined *S. aureus* can attach to oropharyngeal epithelium [37–39], but investigators have not determined the biologic mechanism for this attachment. Thus, future studies should assess mechanisms by which *S. aureus* may attach to oropharyngeal tissue.

## Acknowledgements

The authors would like to thank Dr. Elizabeth Chrischilles and Dr. Kelley Donham for their guidance during the analysis of the study.

